# Ability of known susceptibility SNPs to predict colorectal cancer risk for persons with and without a family history

**DOI:** 10.1101/267666

**Authors:** Mark A. Jenkins, Aung K. Win, James G. Dowty, Robert J. MacInnis, Enes Makalic, Daniel F. Schmidt, Gillian S. Dite, Miroslaw Kapuscinski, Mark Clendenning, Christophe Rosty, Ingrid M. Winship, Jon D. Emery, Sibel Saya, Finlay A. Macrae, Dennis J. Ahnen, David Duggan, Jane C. Figueiredo, Noralane M. Lindor, Robert W. Haile, John D. Potter, Michelle Cotterchio, Steven Gallinger, Polly A. Newcomb, Daniel D. Buchanan, Graham Casey, John L. Hopper

**Author notes:** Corresponding Author: Mark Jenkins, PhD, Centre for Epidemiology and Biostatistics, Melbourne School of Population and Global Health, Level 3, 207 Bouverie Street, The University of Melbourne VIC 3010 Australia.

## Abstract

**Background:** A number of single nucleotide polymorphisms (SNPs), which are common inherited genetic variants, have been identified that are associated with risk of colorectal cancer. The aim of this study was to determine the ability of these SNPs to estimate colorectal cancer (CRC) risk for persons with and without a family history of CRC, and the screening implications.

**Methods:** We estimated the association with CRC of a 45 SNP-based risk using 1,181 cases and 999 controls, and its correlation (*r*) with CRC risk predicted from detailed family history. We estimated the predicted change in the distribution across predefined risk categories, and implications for recommended age to commence screening, from adding SNP-based risk to family history.

**Results:** The inter-quintile risk ratio for colorectal cancer risk of the SNP-based risk was 2.46 (95% CI 1.91 – 3.11). SNP-based and family history-based risks were not correlated (*r* = 0.02). For persons with no first-degree relatives with CRC, recommended screening would commence 2 years earlier for women (4 years for men) in the highest quintile of SNP-based risk, and 12 years later for women (7 years for men) in the lowest quintile. For persons with two first-degree relatives with CRC, recommended screening would commence 15 years earlier for men and women in the highest quintile, and 8 years earlier for men and women in the lowest quintile.

**Conclusions:** Risk reclassification by 45 SNPs could inform targeted screening for CRC prevention, particularly in clinical genetics settings when mutations in high-risk genes cannot be identified.

## INTRODUCTION

Categorisation of people by their colorectal cancer (CRC) risk can guide risk-based prevention, including screening. Family history of the disease is a well-established risk factor for CRC; accordingly screening guidelines recommend that screening be greater for those with a family history compared to those without a family history of the disease.[1] This increased screening could be by modality (e.g. colonoscopy vs faecal occult blood test [FOBT]), age at which screening commences (younger vs older) or frequency (e.g. two yearly vs every 10 years), or a combination of these in equipoise, based on the cost-effective use of limited resources and safety. For persons with a strong family history, efforts to identify the heritable basis of the disease can involve germline mutation screening and testing such as the DNA mismatch repair genes responsible for Lynch syndrome. In this clinical setting, current practice is finding that for a substantial proportion of such families no cancer predisposing mutation can be identified, leaving no other option but to offer screening based on the average risk of CRC based on the cancer family history alone. Family history is a blunt measure of increased risk, as even within groups of persons with the same family history there is substantial variation in the risk of CRC.[2] This suggests the existence of other genetic risk factors which, if identified, could be used to further stratify risk allowing for a more appropriate screening, compared with that recommended based on family history alone.

Recently, genome-wide association studies (GWASs) have found multiple single nucleotide polymorphisms (SNPs) associated with the risk of CRC. Although each SNP is associated with only a small increment in risk, combining these SNPs has the potential to improve risk estimation. The CRC risk based on the first 10 SNPs discovered was sufficient to identify the 0.4% of the population whose risk exceeded the threshold (5% over a 10-year period) for which colonoscopy screening, rather than the less invasive FOBT, would be recommended.[3] Since that report, more independent SNPs associated with CRC risk have been identified, and risk gradients associated with 14 and 27 additional independent SNPs have been published.[4, 5]

In 2015, we published a theoretical evaluation of the 45 independent SNPs identified from a systematic literature review that had been internally or externally validated to be associated with CRC risk for persons of European ancestry.[6] We predicted that the 20% of the population with the highest number of risk alleles would be at 1.8-times the risk of persons with average number of risk alleles. Consequently, this group would be predicted to attain average population age-specific risk approximately 9 years earlier than would someone with the average number of risk alleles. We also predicted that the 45 CRC-risk associated SNPs identified in the literature could explain about 22% of the familial component of CRC risk.[6]

To evaluate this theoretical SNP-based risk, and to determine its clinical utility, we have conducted a validation study using a population-based sample of CRC cases and controls, and assessed its ability to improve risk classification and change recommended CRC screening of people compared with classification based on family history alone.

## MATERIALS AND METHODS

### Study sample

We used CRC cases and controls from the Colon Cancer Family Registry, which has been described in detail previously.[7] Cases were persons with invasive cancer of the colon or rectum identified from population-based cancer registries in the Puget Sound region of the state of Washington in the USA, Ontario in Canada, and Victoria in Australia. Controls were persons who had not had a diagnosis of CRC randomly selected from the general population by using Medicare and Driver’s License files (Washington, USA), telephone subscriber lists (Ontario, Canada), or electoral rolls (Victoria, Australia).

For the estimation of a SNP-based risk, we used 1,181 cases and 999 controls who underwent genome-wide testing from a GWAS using the Illumina Human1M v1 or Illumina Human1M-Duo v3.0 platform. Given that the original purpose of the GWAS was to identify new CRC susceptibility genes, cases were preferentially selected to be aged younger than 50 years at diagnosis with a 10% sampling from all other ages at diagnoses, and controls were preferentially selected to not have a family history of CRC. Cases were tested for germline mutations in the DNA mismatch repair genes and *MUTYH*, and all mutation carriers were excluded. Informed consent was obtained from all study participants, and the study protocol was approved by the institutional research ethics review board at each study site.

### Estimation of SNP-based risk

For each case and control, we estimated an individual SNP-based risk based on the 45 SNPs that we previously selected[6, 8] as being associated with CRC risk from a search of the literature. Using the approach of Mealiffe et al.,[9] we estimated for each of the 45 SNPs the odds ratio (OR) per risk allele and risk allele frequency (*p*) assuming independent and additive risks on the log OR scale. For each SNP, we calculated the population average risk as *μ* = (1 − *p*)^2^ + 2*p*(1 − *p*)OR + *p*^2^OR^2^. Weighted risk values (so that the population average risk was equal to 1) were calculated as 1/*μ*, OR/*μ* and OR^2^/*μ* for the three genotypes defined by number of risk alleles (0, 1, or 2). The overall SNP-based risk for each individual was then calculated by multiplying the weighted risk values for each of the 45 SNPs (Supplement Table 1).

**Table 1.** Characteristics of cases and controls used for estimation of the SNP-based risk of colorectal cancer

|  |  | Cases<br>(n=1,181) |  | Controls<br>(n=999) |  |
| --- | --- | --- | --- | --- | --- |
|  |  | N | (%) | N | (%) |
| Sex |  |  |  |  |  |
|  | Women | 567 | (48.0) | 521 | (52.1) |
|  | Men | 614 | (52.0) | 478 | (47.9) |
| Country |  |  |  |  |  |
|  | Canada | 417 | (35.3) | 501 | (50.2) |
|  | Australia | 320 | (27.1) | 189 | (18.9) |
|  | USA | 444 | (37.6) | 309 | (30.9) |
| Age (years) |  |  |  |  |  |
|  | <40 | 111 | (9.4) | 48 | (4.8) |
|  | 40-49 | 450 | (38.1) | 113 | (11.3) |
|  | 50-59 | 263 | (22.3) | 304 | (30.4) |
|  | 60-69 | 232 | (19.6) | 305 | (30.5) |
|  | ≥70 | 125 | (10.6) | 229 | (22.9) |
|  | Mean (SD), years | 53.0 | (11.4) | 59.9 | (11.0) |
| Risk alleles* |  |  |  |  |  |
|  | Mean (SD) | 42.7 | (4.3) | 40.9 | (4.4) |
|  | Median (range) | 42 | (30-57) | 41 | (25-54) |

\* possible range is 0 – 90 (two alleles for each of the 45 SNPs)

The SNP-based risk was log transformed for all analyses. We estimated the association between SNP-based risk and CRC by applying multiple logistic regression to the case and control data. We adjusted for age group, sex and recruitment site.

We assessed the risk gradient, and hence the discrimination in risk between cases and controls, by estimating the change in odds per adjusted standard deviation (OPERA). OPERA interprets risk estimates by adjusting the standard deviation for the other factors taken into account by design and analysis.[10] We also estimated risk discrimination by the inter-quintile risk ratio (average CRC risk for those in the top 20% of the population for the SNP score divided by average risk for those in the bottom 20% of the population for the SNP score). The inter-quintile risk-ratio was derived by exponentiating the OPERA estimate by 2.8. Given the deliberate deficit of controls with a family history, the estimated SNP-based association was adjusted down by 4% based on the theoretical gradient of polygenic risk in the population (see details in Supplement).[11] For prediction in terms of age- and sex-specific population incidences, we used data from the Surveillance, Epidemiology, and End Results Program Cancer Statistics.[12]

We assessed the extent to which the SNPs were dependent on familial risk by estimating the Pearson’s correlation (*r*) between the family history-based and SNP-based risks using the cases in the GWAS dataset. For the estimation of family history-based risk, we calculated the lifetime absolute risk (probability) of CRC predicted by a mixed major gene–polygenic model.[13] The model estimates each person’s risk of CRC using detailed family history data. It considers, for each relative, the age at diagnosis of CRC as well as their relationship to the proband, age at last living or age at death, and their high-risk gene mutation status, if known.

We also assessed the ability to reclassify the recommended age at commencement of screening by including the SNP-based risk. The 5-year risk of CRC for the average person (without a previous diagnosis of CRC) in the USA is approximately 0.3% at age 50 years,[14] which is the age that guidelines recommend screening to commence in many countries including the USA.[15] We estimated the ages at which the average woman and man in the highest and lowest quintile of SNP-based risk met this 0.3% risk threshold.

Stata version 14.2[16] was used for all statistical analyses unless otherwise specified. All statistical tests were two-sided, and *P*-values less than 0.05 were considered nominally statistically significant. Detailed statistical methods for the calculation of risk distributions are provided in the Supplement.

## RESULTS

Cases and controls were balanced for sex, cases were distributed almost evenly across participating regions, controls were predominantly from Canada, and cases had on average 1.8 more risk alleles than the controls (Table 1). The OR per adjusted standard deviation for SNP-based risk was 1.38 (95% CI 1.26 – 1.50, p <0.001). The corresponding inter-quintile risk ratio was 2.46 (1.91 – 3.11). The correlation between the SNP-based risk and the family history-based risk was *r* = 0.02.

Figure 1 shows the distribution of lifetime risk of CRC to age 80 years for the US population by SNP-based risk and family history categories. Persons with no first-degree relative with CRC constitute 90% of the population and have an average lifetime CRC risk to age 80 years of 4.0%, which is 10% lower than the population average risk of 4.4%. Of persons with no first-degree relative with CRC, those in the highest quintile for SNP-based risk have an average risk of 6.0%, (~50% higher than those with the average SNP-based risk). Of persons with no first-degree relative with CRC and in the highest quintile for SNP-based risk, approximately 1 in 3 (32%) have a ‘low’ CRC risk (lifetime risk less than 2%) and approximately 1 in 50 (2%) have a ‘very high’ CRC risk (lifetime risk of 30% or greater). Those with no first-degree family history and in the lowest quintile for SNP-based risk have an average risk of 3%. Of these, most (61%) have a ‘low’ CRC risk and only 1 in 500 (0.2%) have a ‘very high’ risk (Table 2 and Figure 2).

**Figure 1.**
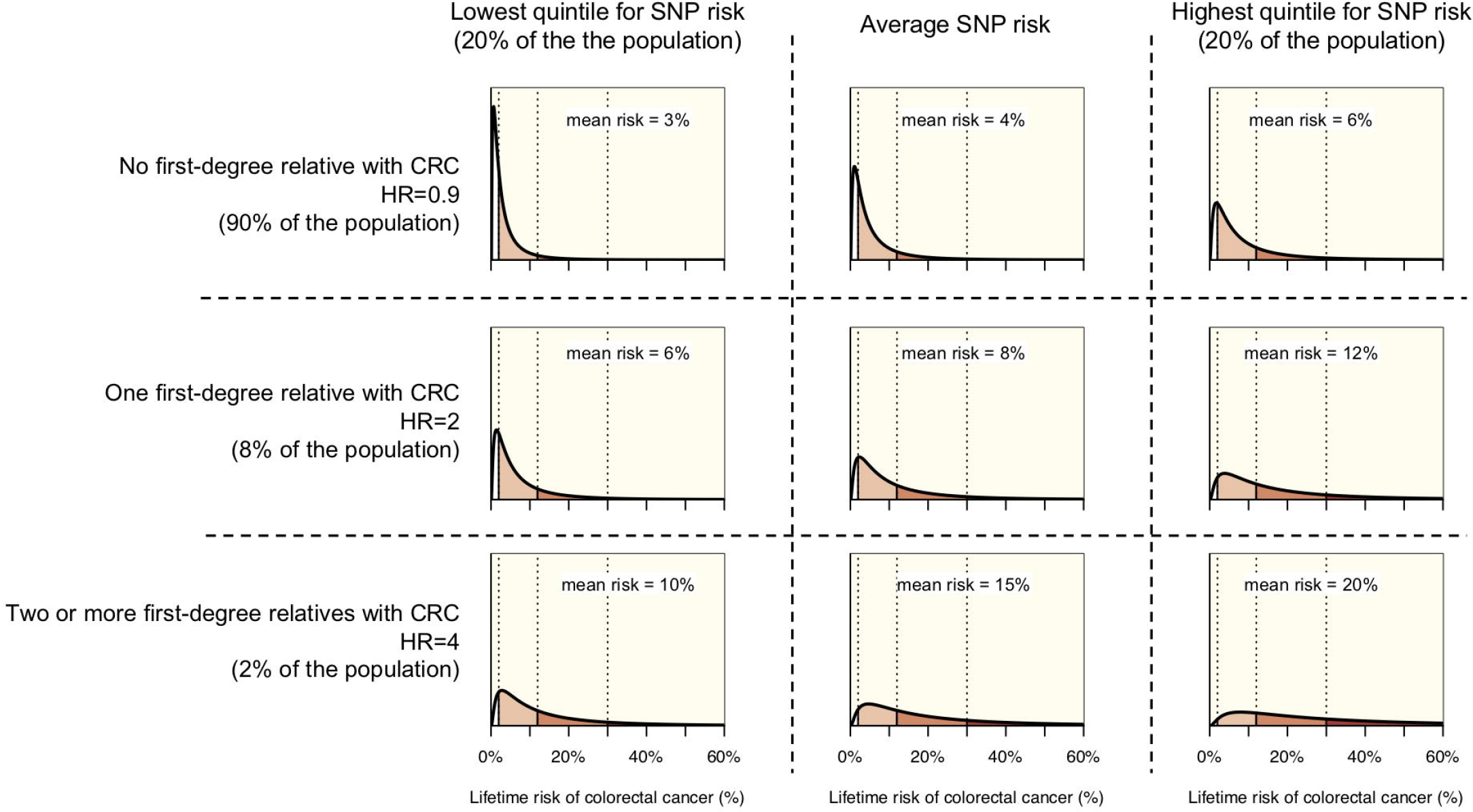
The distribution of lifetime risk of colorectal cancer (CRC) i.e., cumulative risk to age 80 years for the US population, by three categories of SNP-based risk and three categories of family history. Risks are shown for those in the lowest quintile of SNP-based risk (left column), those at average risk (centre column), and those in the highest quintile of SNP-based risk (right column) by for those with no first-degree relative with CRC (top row), those with one first-degree relative with CRC (middle row), and those with two or more first-degree relatives with CRC (bottom row).

**Table 2.**
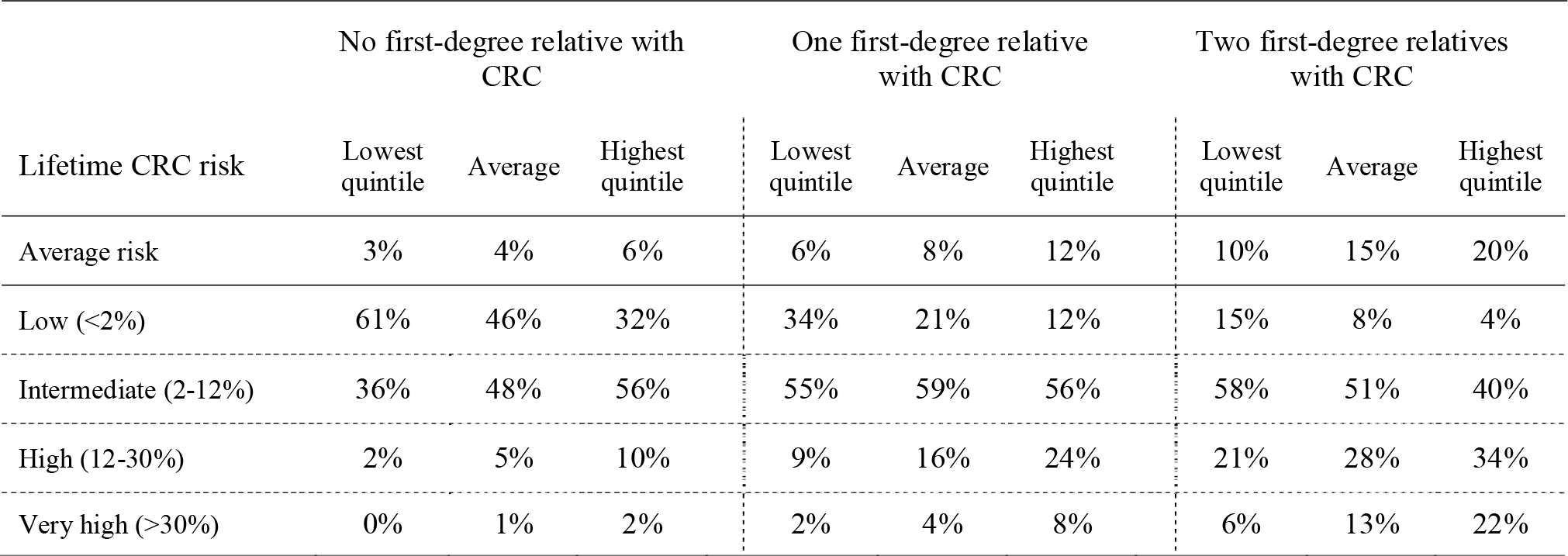
The distribution of lifetime risk of colorectal cancer (CRC) i.e., cumulative risk to age 80 years for the US population, by categories of family history of CRC, separately for persons in the lowest quintile, average, and highest quintile of SNP-based risk.

**Figure 2.**
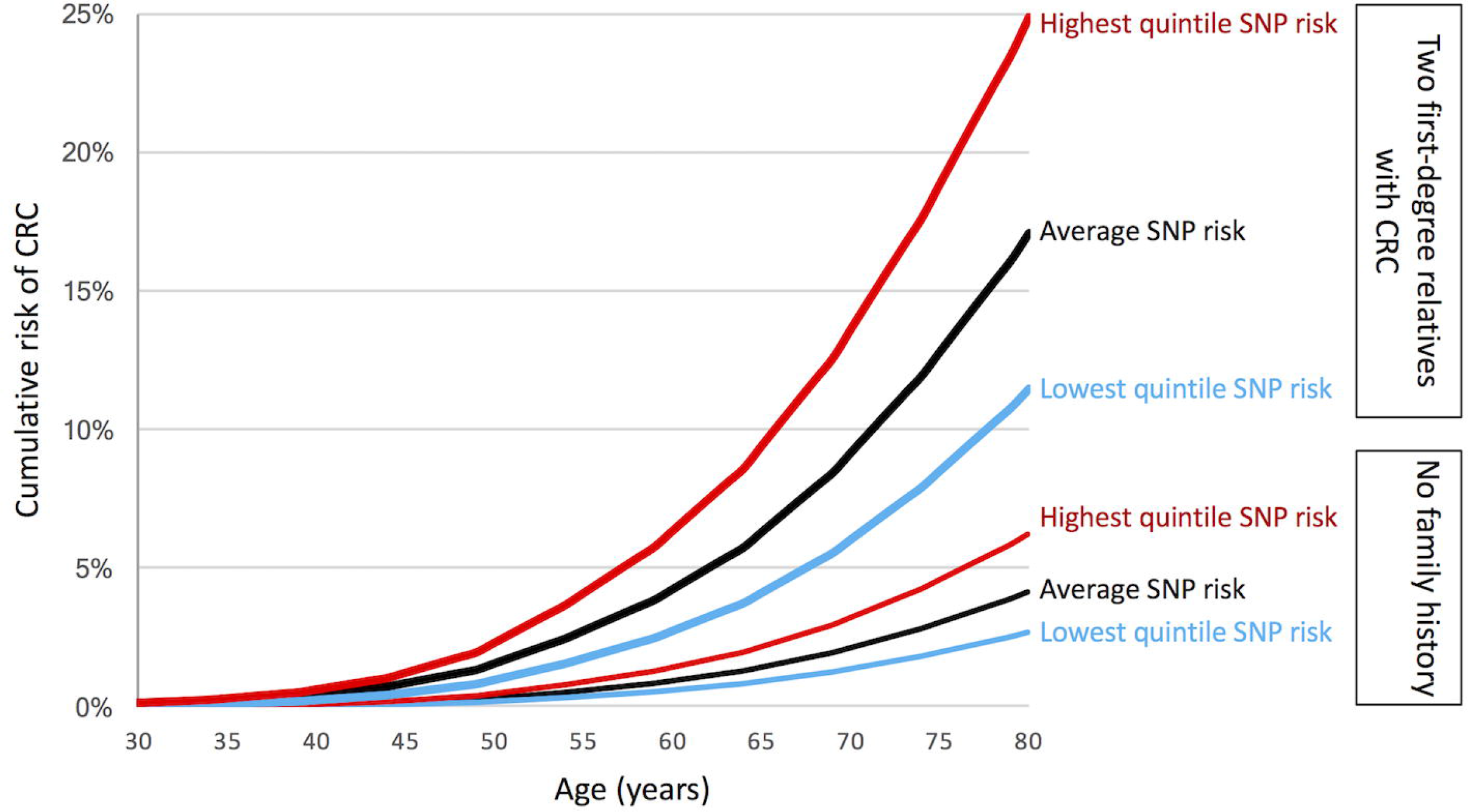
Cumulative risk of colorectal cancer (CRC) for the US population by three categories of SNP-based risk (lowest quintile, average, highest quintile) and two categories of family history (no first-degree relatives with CRC labelled “No family history” and two first-degree relatives with CRC).

At the other end of the scale of family history, 2% of the US population have two or more first-degree relatives with CRC, with an average lifetime CRC risk of 4-times that for the general population. For persons with two first-degree relatives with CRC, the average risk is 15%. But for those in the highest quintile for SNP-based risk, the average risk is 20%. Of these, only 1 in 25 (4%) have a ‘low’ CRC risk while approximately one in five (22%) have a ‘very high’ CRC risk. In contrast, even though they have a strong family history, those in the lowest quintile of SNP-based risk have an average risk of 10%, with only one in 6 (15%) having a ‘low’ risk and one in 15 (6%) having a ‘very high’ risk (Table 2 and Figure 2).

The addition of a SNP-based risk to the family history-based risk identified persons younger than age 50 who had a CRC risk at least as high as an average 50-year-old which is approximately 0.3% (Figure 2). For women with no first-degree relatives with CRC and in the highest quintile for SNP-based risk, their CRC risk is at least this high by the age of 48 (i.e. 2 years younger), with a corresponding age for men at 46 years (4 years earlier). For those with first-degree relatives with CRC and in lowest quintile of SNP-based risk, this risk threshold is not reached until 62 years for women and 57 years for men. For those with two-first degree relatives and being in the highest quintile for SNP-based risk, the 0.3% CRC risk is achieved at age 35 years for women and men (i.e. 15 years younger), and in those in the lowest quintile for risk, the threshold is reached at ages 42 years for both women and men. (Figure 2 and Table 3).

**Table 3.** Age (years) at which a person’s 5-year risk of colorectal cancer (CRC) reaches or exceeds the threshold of 0.3%, by quintile status of the 45 SNP-based risk and categories of family history of CRC, separately for women and men.

| Family history | SNP-based risk | Women | Men |
| --- | --- | --- | --- |
| All | Highest quintile | 47 | 45 |
|  | Lowest quintile | 61 | 55 |
| No first-degree relatives with CRC | Highest quintile | 48 | 46 |
|  | Lowest quintile | 62 | 57 |
| One first-degree relative with CRC | Highest quintile | 41 | 40 |
|  | Lowest quintile | 48 | 47 |
| Two first-degree relatives with CRC | Highest quintile | 35 | 35 |
|  | Lowest quintile | 42 | 42 |

## DISCUSSION

CRC risk estimation is important because screening can result in prevention, or at least early intervention, through early detection and removal of precancerous lesions and CRC, with more effective treatment of CRC at an earlier stage. Screening comprises different modalities, including FOBT and colonoscopy, and use should be tailored to the level of personal risk. FOBT is inexpensive and safe, but may be less sensitive for pre-cancerous polyps; while colonoscopy is expensive, invasive and carries its own risks (bleeding, perforation and infrequently even death). We have shown that risk information from 45 independent SNPs adds to the risk information from using family history alone. Importantly, on an individual level, SNP-based risk estimation can lead to a risk category that is higher or lower than when estimated solely through family history.

Based on a population of the same size and sex- and age-distribution, and the same sex- and age-specific CRC incidence as the United States, we have estimated the number people who would screen earlier and later if their SNP-based risk was used to determine screening starting age (as provided in Table 4). If screening starting age was based on SNP-based risk, we estimate that 3.32 million people aged 46 to 50 years would be in the highest quintile of SNP-based risk and therefore could begin screening at age 46 (4 years younger than the general population). We estimate that each year approximately 8,000 of these people would be diagnosed with CRC i.e., cancers that would not have been screened for if SNP-based risk was not used to guide screening start age. This equates to screening 415 people for every CRC. We also estimate that 8.76 million people aged between 50 and 59 years would be in the lowest quintile of SNP-based risk and therefore, if SNP scores were used to decide which people would delay screening, would be 9 years older than the general population when they begin screening. We estimate in the 9 years from age 50 to 59, approximately 18,000 of these people would be diagnosed with CRC i.e., cancers that would not been screened for if SNP-based risk was used to guide screening start age. If these were screened, it would equate to screening 486 people for every CRC. Therefore, basing screening starting age on SNP-based risk would result in an overall increase in efficiency for CRC detection and identify a new group of at-risk people under 50 for whom screening could be justified. However delaying screening for the lowest quintile for SNP-based risk would miss a screening opportunity when many CRCs would be occurring.

For the vast majority of the population who have no first-degree family history of CRC, many guidelines recommend that screening should commence at age 50 years (commonly by biennial FOBT or 10-yearly colonoscopy). Our analyses suggest that the SNP-based risk, especially when combined with family history, can identify subsets that would be recommended to have a higher level of screening, perhaps starting at a younger age. Of the 90% of the population with no parent, sibling or child with CRC, use of SNPs could identify the 20% of these with the highest SNP-based risk who have an average CRC risk of 6%, which is 50% higher than the average risk and screening commencement would be recommended to commence 2 to 4 years early. Further, of these 20% with the highest SNP-based risk, 12% have a high or very high risk of CRC compared with only 6% of those with an average SNP-based risk, i.e. SNP-based risk assessment can result in a 2-fold enrichment of CRC risk even in those with no first-degree relatives with CRC.

For persons with a family history of CRC, many guidelines recommend an increased level of screening (e.g. five-yearly colonoscopy). These people are often referred to familial cancer clinics for genetic screening for mutations in major CRC susceptibility genes including the DNA mismatch repair genes. If identified, high-risk gene carriers can be offered a higher level of screening (e.g., annual colonoscopy). Unfortunately, due in part to the rarity of mutation carriers in these genes (even for persons with a family history), in current practice a mutation cannot be identified in the majority of patients screened because they have a family history. Our analyses show that, within family history categories, SNP-based risk assessment can identify persons who belong in lower or higher risk categories.

For persons with a strong family history, such as two first-degree relatives with CRC, the ability of the SNP-based risk assessment to reclassify risk is even more apparent. Those in top 20% for SNP-based risk have an average CRC risk of 20%, and would be recommended to commence screening 15 years earlier. More than half of them will be at high or very high CRC risk. With the same strong family history, those 20% with the lowest SNP-based risk had an average CRC risk of 10%, and they would be recommended to commence screening 8 years earlier. Just over one quarter of them will be at high or very high CRC risk.

We found that the CRC risk based on 45 SNPs was not appreciably correlated with the family history-based risk, which means the increased risks due to family history and SNPs are virtually independent and their associations are likely multiplicative (as has been found for breast cancer[17]). Therefore, both are important risk factors to consider in order to estimate CRC risk. This also means that the 45 SNPs explain little of the reason why CRC aggregates in families, or why CRC in a relative is associated with an increased risk of CRC.

A potential limitation of our study is that we used a case-control dataset in which the controls were selected for not having a family history. We therefore had to reduce the observed SNP associations by 4% to account for the controls being over-sampled for not having a family history.

There could be many more yet to be discovered independent SNPs associated with the risk of CRC, and there could be interactions between SNPs within and across different genes.[18, 19] A SNP-based risk prediction model is likely to perform better when these SNPs are discovered, for example by using larger sample sizes or by fine mapping genomic regions of interest such as those identified by novel approaches such as DEPTH,[20] and included in risk prediction models. Analytic approaches that extract more information from genotyping data by, for example, using machine learning to consider all SNPs that lead to an improvement in risk prediction,[21] especially by focusing on pathways or SNP–SNP interactions, might also produce better SNP-based risk prediction.

To explain the on average 2-fold increased CRC risk associated with having one first-degree relative with CRC, mathematical models predict that the familial component of CRC risk must have a very large variance, so large that the risk for persons in the upper quartile would be at least 20-times than the risk for persons in the lower quartile.[11] This study has shown that, by using risk information based on both SNPs and detailed family history, a non-trivial proportion of this variance can being explained, and we have quantified the ability to differentiate between persons at low risk (much less than population average risk) and those at increased risk, across a very wide range. Given that CRC can be effectively prevented by screening, and mortality from the disease reduced by early detection, risk assessment based on SNPs together with other risk factors including family history enables the possibility of precision prevention and screening to substantially lower the impact of CRC.[2]

Our modelling exercise only considers family history and SNP-based risk as we have focussed this paper on inherited risks. However, other factors do contribute to CRC risk (e.g. lifestyle factors) and a full risk-based assessment for screening could include these factors in addition to family history and SNPs which would result in a greater risk discrimination.[22]

If new guidelines on screening were to adopt SNP-based risk assessment, our study suggests that the screening guidelines for CRC would be substantially altered: (a) For those with 2 first-degree relatives with CRC, screening would commence at age 35 years for both women and men in the highest quintile for SNP-based risk; and at age 42 years for both women and men in the lowest quintile; (b) For those with one first-degree relatives with CRC, screening would commence at age 40 years for both women and men in the highest quintile of SNP-based risk, and at age 47 years for both women and men in the lowest quintile; and (c) For those with no first-degree relatives with CRC, screening would commence at age 48 years for women and at age 46 years for men in the highest quintile, and at age 62 years for women and at age 57 years for men in the lowest quintile.

While this is an important first step, we agree that many issues would need to be resolved before SNP-based risk was incorporated into standard of care for CRC screening, that are beyond the scope of this study. These include assessment of cost-effectiveness, resources requirements, community, patient and clinician acceptance and feasibility of incorporation within existing screening programs, with potentially ethical, legal and insurance implications. Cost-effectiveness implications of this research are indeed important for screening programs, especially as a new personalised risk-based approach to screening is designed to optimise risk vs benefit compared with conventional approaches to (moderate/high) risk and colonoscopy. Any improvement as promised with the current approach is an important contribution to public health and risk management.

In conclusion, we have shown that risk information from considering the 45 SNPs and a detailed family history together can result in substantial reclassification of risk category for various levels of family history, including those without a family history. It is therefore important to include *both* family history and SNP assessment when estimating CRC risk. This new risk measure could inform targeted screening and prevention, for the general population young than age 50, and in the clinical setting for those in whom a high-risk gene mutation cannot be identified.

## Declarations

### Ethics approval and consent to participate

Informed consent was obtained from all study participants, and the study protocol was approved by the institutional research ethics review board at each study site.

### Consent for publication

Not applicable

### Availability of data and material

The datasets used and/or analysed during the current study are available from the corresponding author on reasonable request.

### Authors’ contributions

MAJ conceived the aims; MAJ, AKW, JGD, RJM conducted the analyses and were major contributors to the manuscript; EM, DFS, GSD, MK derived the SNP scores and provided contributions to the manuscript; CR, IMW, JDE, SS, FAM, DJA, DD provided interpretation of the findings and provided contributions to the manuscript; JCF, NML, RWH, JDP, MC, SG, PAN, DDB, GC, JLH led recruitment and genetic testing and provided contributions to the manuscript. All authors read and approved the final manuscript.

## Acknowledgements

The authors thank all study participants of the Colon Cancer Family Registry and staff for their contributions to this project.

## DISCLAIMER

The content of this manuscript does not necessarily reflect the views or policies of the National Cancer Institute or any of the collaborating centres in the Colon Cancer Family Registry, nor does mention of trade names, commercial products, or organizations imply endorsement by the US Government or the Colon Cancer Family Registry. Authors had full responsibility for the design of the study, the collection of the data, the analysis and interpretation of the data, the decision to submit the manuscript for publication, and the writing of the manuscript.

## REFERENCES

1. Win AK, Ait Ouakrim D, Jenkins MA: Risk profiling: familial colorectal cancer. Cancer Forum 2014, 38(1):15–25.

2. Hopper JL: Disease-specific prospective family study cohorts enriched for familial risk. Epidemiol Perspect Innov 2011, 8(1):2.

3. Dunlop MG, Tenesa A, Farrington SM, Ballereau S, Brewster DH, Kossler T, Pharoah P, Schafmayer C, Hampe J, Volzke H et al.: Cumulative impact of common genetic variants and other risk factors on colorectal cancer risk in 42103 individuals. Gut 2013, 62(6):871–881.

4. Lubbe SJ, Di Bernardo MC, Broderick P, Chandler I, Houlston RS: Comprehensive evaluation of the impact of 14 genetic variants on colorectal cancer phenotype and risk. American journal of epidemiology 2012, 175(1):1–10.

5. Hsu L, Jeon J, Brenner H, Gruber SB, Schoen RE, Berndt SI, Chan AT, Chang-Claude J, Du M, Gong J et al.: A model to determine colorectal cancer risk using common genetic susceptibility Loci. Gastroenterology 2015, 148(7):1330–1339 e1314.

6. Jenkins MA, Makalic E, Dowty JG, Schmidt DF, Dite GS, MacInnis RJ, Ait Ouakrim D, Clendenning M, Flander LB, Stanesby OK et al.: Quantifying the utility of single nucleotide polymorphisms to guide colorectal cancer screening. Future Oncol 2016, 12(4):503–513.

7. Newcomb PA, Baron J, Cotterchio M, Gallinger S, Grove J, Haile R, Hall D, Hopper JL, Jass J, Le Marchand L et al.: Colon Cancer Family Registry: an international resource for studies of the genetic epidemiology of colon cancer. Cancer Epidemiol Biomarkers Prev 2007, 16(11):2331–2343.

8. Stanesby O, Jenkins M: Comparison of the efficiency of colorectal cancer screening programs based on age and genetic risk for reduction of colorectal cancer mortality. Eur J Hum Genet 2017, 25(7):832–838.

9. Mealiffe ME, Stokowski RP, Rhees BK, Prentice RL, Pettinger M, Hinds DA: Assessment of clinical validity of a breast cancer risk model combining genetic and clinical information. J Natl Cancer Inst 2010, 102(21):1618–1627.

10. Hopper JL: Odds per Adjusted Standard Deviation: Comparing Strengths of Associations for Risk Factors Measured on Different Scales and Across Diseases and Populations. Am J Epidemiol 2015, 182(10):863–867.

11. Hopper JL, Carlin JB: Familial Aggregation of a Disease Consequent upon Correlation between Relatives in a Risk Factor Measured on a Continuous Scale. Am J Epidemiol 1992, 136(9):1138–1147.

12. Howlader N NA, Krapcho M, Garshell J, Miller D, Altekruse SF, Kosary CL, Yu M, Ruhl J, Tatalovich Z,Mariotto A, Lewis DR, Chen HS, Feuer EJ, Cronin KA (eds). SEER Cancer Statistics Review, 1975-2011,External Web Site IconNational Cancer Institute. Bethesda, MD, http://seer.cancer.gov/csr/1975_2011/browse_csr.php?sectionSEL=6&pageSEL=sect_06_table.10.html based on November 2013 SEER data submission, posted to the SEER Web site, April 2014.

13. Win AK, Jenkins MA, Dowty JG, Antoniou AC, Lee A, Giles GG, Buchanan DD, Clendenning M, Rosty C, Ahnen DJ et al.: Prevalence and Penetrance of Major Genes and Polygenes for Colorectal Cancer. Cancer epidemiology, biomarkers & prevention: a publication of the American Association for Cancer Research, cosponsored by the American Society of Preventive Oncology 2017, 26(3):404–412.

14. Howlader N, Noone A, Krapcho M, Garshell J, Miller D, Altekruse S, Kosary C, Yu M, Ruhl J, Tatalovich Z et al. (eds.): SEER Cancer Statistics Review, 1975-2011. Bethesda, MD: National Cancer Institute; 2014.

15. Force USPST, Bibbins-Domingo K, Grossman DC, Curry SJ, Davidson KW, Epling JW, Jr., Garcia FAR, Gillman MW, Harper DM, Kemper AR et al.: Screening for Colorectal Cancer: US Preventive Services Task Force Recommendation Statement. Jama 2016, 315(23):2564–2575.

16. StataCorp: Stata Statistical Software: Release 14. In. College Station, TX: StataCorp LP; 2015.

17. Dite GS, MacInnis RJ, Bickerstaffe A, Dowty JG, Allman R, Apicella C, Milne RL, Tsimiklis H, Phillips KA, Giles GG et al.: Breast Cancer Risk Prediction Using Clinical Models and 77 Independent Risk-Associated SNPs for Women Aged Under 50 Years: Australian Breast Cancer Family Registry. Cancer epidemiology, biomarkers & prevention: a publication of the American Association for Cancer Research, cosponsored by the American Society of Preventive Oncology 2016, 25(2):359–365.

18. Frampton MJ, Law P, Litchfield K, Morris EJ, Kerr D, Turnbull C, Tomlinson IP, Houlston RS: Implications of polygenic risk for personalised colorectal cancer screening. Ann Oncol 2016, 27(3):429–434.

19. Tenesa A, Dunlop MG: New insights into the aetiology of colorectal cancer from genome-wide association studies. Nat Rev Genet 2009, 10(6):353–358.

20. MacInnis RJ, Schmidt DF, Makalic E, Severi G, FitzGerald LM, Reumann M, Kapuscinski MK, Kowalczyk A, Zhou Z, Goudey B et al.: Use of a Novel Nonparametric Version of DEPTH to Identify Genomic Regions Associated with Prostate Cancer Risk. Cancer Epidemiol Biomarkers Prev 2016, 25(12):1619–1624.

21. Wei Z, Wang W, Bradfield J, Li J, Cardinale C, Frackelton E, Kim C, Mentch F, Van Steen K, Visscher PM et al.: Large sample size, wide variant spectrum, and advanced machine-learning technique boost risk prediction for inflammatory bowel disease. Am J Hum Genet 2013, 92(6):1008–1012.

22. Frampton M, Houlston RS: Modeling the prevention of colorectal cancer from the combined impact of host and behavioral risk factors. Genet Med 2017, 19(3):314–321.

